# Information Confusion Reveals an Innate Limit of the Information Processing by Neurons

**DOI:** 10.1101/2020.10.19.345249

**Authors:** Yang Tian, Justin L. Gardner, Guoqi Li, Pei Sun

## Abstract

Information experiences complex transformation processes in the brain, involving various errors. A daunting and critical challenge in neuroscience is to understand the origin of these errors and their effects on neural information processing. While previous efforts have made substantial progresses in studying the information errors in bounded, unreliable and noisy transformation cases, it still remains elusive whether the neural system is inherently error-free under an ideal and noise-free condition. This work brings the controversy to an end with a negative answer. We propose a novel neural information confusion theory, indicating the widespread presence of information confusion phenomenon after the end of transmission process, which originates from innate neuron characteristics rather than external noises. Then, we reformulate the definition of zero-error capacity under the context of neuroscience, presenting an optimal upper bound of the zero-error transformation rates determined by the tuning properties of neurons. By applying this theory to neural coding analysis, we unveil the multi-dimensional impacts of information confusion on neural coding. Although it reduces the variability of neural responses and limits mutual information, it controls the stimulus-irrelevant neural activities and improves the interpretability of neural responses based on stimuli. Together, the present study discovers an inherent and ubiquitous precision limitation of neural information transformation, which shapes the coding process by neural ensembles. These discoveries reveal that the neural system is intrinsically error-prone in information processing even in the most ideal cases.

**Author summary:** One of the most central challenges in neuroscience is to understand the information processing capacity of the neural system. Decades of efforts have identified various errors in nonideal neural information processing cases, indicating that the neural system is not optimal in information processing because of the widespread presences of external noises and limitations. These incredible progresses, however, can not address the problem about whether the neural system is essentially error-free and optimal under ideal information processing conditions, leading to extensive controversies in neuroscience. Our work brings this well-known controversy to an end with a negative answer. We demonstrate that the neural system is intrinsically error-prone in information processing even in the most ideal cases, challenging the conventional ideas about the superior neural information processing capacity. We further indicate that the neural coding process is shaped by this innate limit, revealing how the characteristics of neural information functions and further cognitive functions are determined by the inherent limitation of the neural system.

## Introduction

Human brain features the capacity to process the external world information [1, 2]. This information processing process is triggered by external inputs and consists of various forms of information coding in neural ensembles [3–5]. As shown by vast amount of neuroscience studies, neural coding begins with the neural response initiation that is characterized by neural tuning properties [6–8]. Then, the coding process is essentially involves with the spike propagation in neural clusters [9–12], creating ubiquitous information transformation between neurons. Given with the extensive presence of the inter-neuron information transformation, a pivotal and meaningful question is how precisely the information can be transmitted.

Considering the precision limit of neural information transformation, the precision reductions implied by bounded, unreliable and noisy transformation processes are taken into account naturally. It has been unveiled that a significant loss of information is implied when the information amount exceeds the channel capacity of the synapse [13–18], leading to a precision reduction by incomplete information. Moreover, various information errors (e.g., errors caused by the channel unreliability [19] or external random noises [20–22]) are discovered during the transformation, reflecting the fact that precision is frequently limited by the nonideal transformation environments in neural systems. These progresses, however, might lead to a misconception that the precision limit only exists when the transformation is nonideal. Till now, it still lacks of evidence to characterize the neural system as an inherently error-free information processing system.

In this work, we put an end to this controversy by indicating the widespread presence of a kind of innate and noise-independent error, namely information confusion. The confusion happens when different information cases elicit the same receiver response. Different from the information losses and errors during the nonideal transformation, the confusion originates after the transmitted information arrives at the receiver and does not rely on external noises or boundaries. We demonstrate that neural information confusion is caused by the interactions between synaptic connection states and neural tuning properties. Therefore, the precision limit implied by information confusion is intrinsically determined by elementary attributes of the neural system.

Inspired by Shannon’s information theory, we approach this precision limit in the concept of the zero-error capacity, which is the upper limit or supremum of information transfer rates without error (confusion) in a given channel [23–25]. Specifically, we abstract the information space of any given neuron as a graph, which contains all the possible information cases that can be transmitted to this neuron and all the confusion relationships between these cases. Based on previous studies [26, 27], we propose an optimal method to measure the neural zero-error capacity of any neural information space. This systematic theory is demonstrated as mathematically justified and experimentally practical in our work.

We then apply our theory to analyze the effects of information confusion on neural coding. We develop a practical detection method for the information confusion during the coding process. To our surprise, the analysis shows that although the confusion reduces the neural response variability and limits mutual information, it is not completely detrimental for neural coding. This finding may rely on the fact that the stimulus-irrelevant neural activities are hard to survive through the confusion process. These experimental discoveries may deepen the understanding of how neural coding is essentially shaped by the innate neural information processing limit.

## Results

### Neural information confusion

#### Neural information channel

Using a leaky integrate-and-fire model [7, 8, 28], we can simulate the electrodynamics of a neural population (see the section “Leaky integrate-and-fire networks” in Methods). Then we present a model to define the channel between an arbitrary neuron *N*_*i*_ and its receptive field 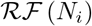.

In neural systems, the intra-system receptive field for a given neuron is defined as the set of all its pre-synaptic neurons. At any moment, each of those pre-synaptic neurons has its own membrane potential state, and there would be an action potential if the membrane potential reaches the spiking threshold. For the given neuron, those states can determine its pre-synaptic inputs, so it is reasonable to treat the ensemble of them as a message sent from the receptive field to this neuron. In this process, the channel is defined as the set of all synaptic connections between the given neuron and its pre-synaptic neurons (**Fig. 1a**).

**Fig 1.**
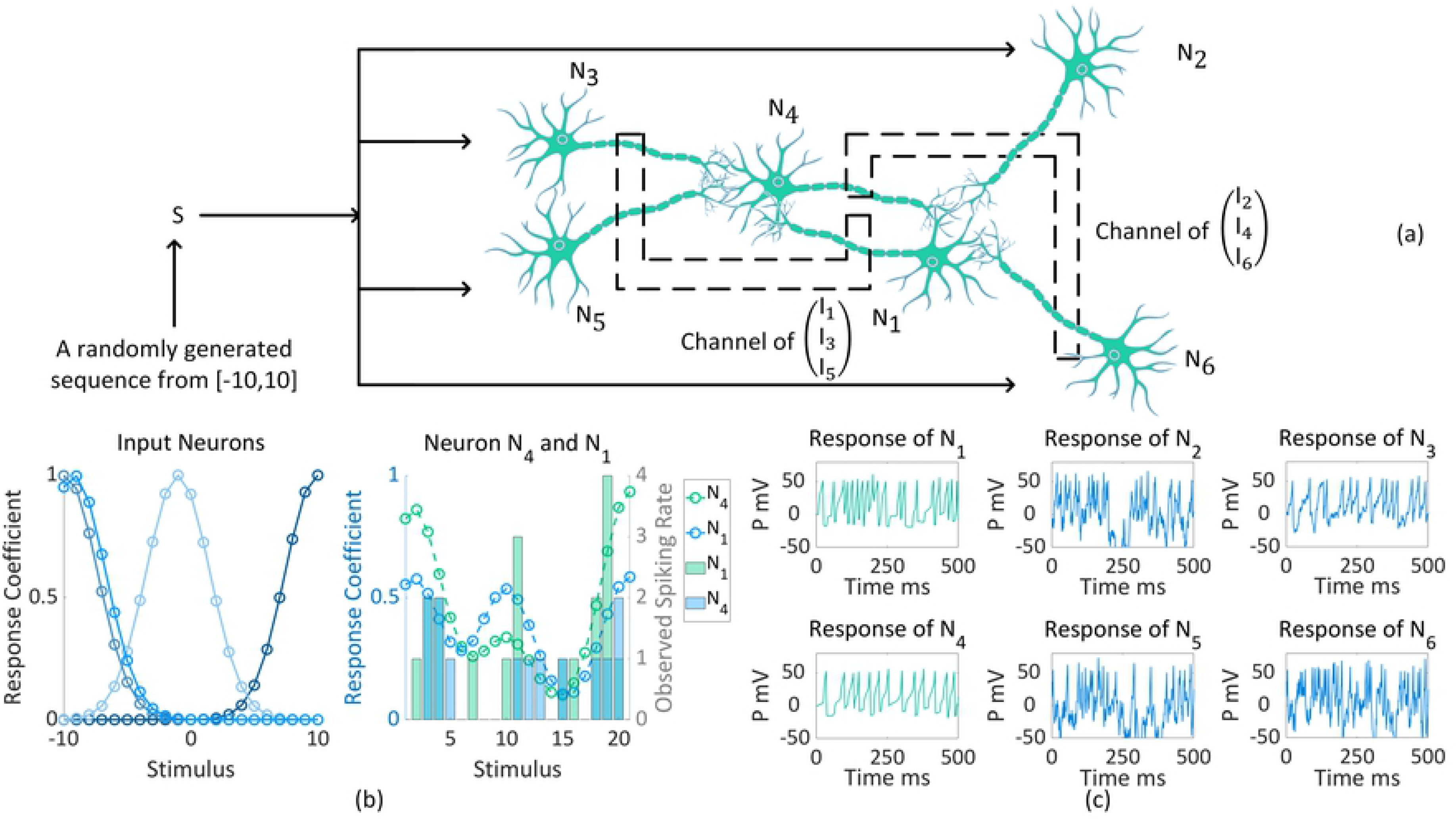
An example of the defined neural population. (**a**) A neural population defined with 6 neurons is set to code the stimulus sequence. The network includes 4 input neurons (*N*_2_, *N*_3_, *N*_5_ and *N*_6_) and 2 intermediary neurons (*N*_1_ and *N*_4_). There are 2 channels showed by dashed boxes in the network, which respectively describe the neural information transformation from {*N*_1_, *N*_3_, *N*_5_} to *N*_4_ as well as that from {*N*_2_, *N*_4_, *N*_6_} to *N*_1_. (**b**) The tuning curve 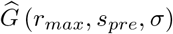 of each input neuron is defined with *s*_*pre*_ ∈ [−10, 10] and 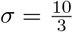. The estimated tuning curves of intermediary neurons (dashed lines with circle markers) share similar variation trends with the observed neural responses (marked by bars). And all tuning curves are shown in the normalized form (*r*_*max*_ = 1). (**c**) The neural responses of the neural population in the interval of 500 ms.

In a neural population, the stimulus inputs may not be received by all neurons at the same time. We refer the neurons that receive stimuli directly as input neurons. For these neurons triggered by other neurons in their receptive fields but not receive stimuli directly, we refer them as intermediary neurons. The response preference of each input neuron is characterized by a bell-shaped tuning curve 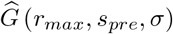, where *r*_*max*_ is the maximum response rate, *s*_*pre*_ is the preferred stimulus and *σ* represents the width of the tuning curve. As for intermediary neurons, we do not define explicitly their activities by presetting their tuning curves since they are not directly triggered by stimuli. Theoretically, we can estimate their tuning properties based on the tuning curves of the neurons in its receptive field and the synaptic connections between them (**Fig. 1b** and the section “Tuning curve and neural response” in Methods). In **Fig. 1b**, it can be seen that the estimated tuning curves share similar trends with the observed neural responses in our experiment, which accords with the common interpretation that neurons tend to make stronger responses at the peaks of tuning curves [6, 29–31]. With a given tuning curve, each neuron *N*_*i*_ can response to stimuli 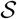 (**Fig. 1c**).

#### Neural information confusion in the information space

The neural information transmits in discrete and finite form. We define the neural information space of neuron *N*_*i*_ as 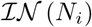, which contains all possible variant cases of the neural information that *N*_*i*_ can receive.

To give a clear vision, we mainly present our theory on the spike-based neural information [32, 33], where the neural response is either 1 for spiking or 0 for non-spiking (we also demonstrate our theory can be applied on the potential-based neural information [34, 35] in the section “Spike-based information space” in Methods). In the neural network, the neural responses of one neuron can also be the bases of synaptic inputs of its post-synaptic neurons. More specifically, in our simplification, the transmitted spike-based information is the excitatory or inhibitory postsynaptic potential. To simulate it, we define the binary vector 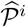 for the neural spiking states of neurons in 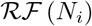 (*N*_*i*_), and the non-recurrent connection strength matrix *C* that indicates the synaptic connections between *N*_*i*_ and other neurons in the neural network (see the section “Spike-based information space” in Methods). Then, let *C*_*i*_ be the *i* column of *C*, the neural information can be represented by the Hadamard product ⊙ of 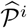 and *C*_*i*_ (**Fig. 2a** and the section “Spike-based information space” in Methods). Thus, for each given neuron *N*_*i*_, the neural information space 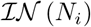 (see the section “Spike-based information space” in Methods, equation (12)) is defined as 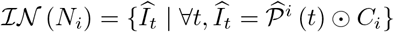, where the matrix *C* can be set as either constant for simplification or dynamical to fit into the plasticity mechanisms (see the section “Neural zero-error capacity definition for the dynamical transformation process” in Methods).

**Fig 2.**
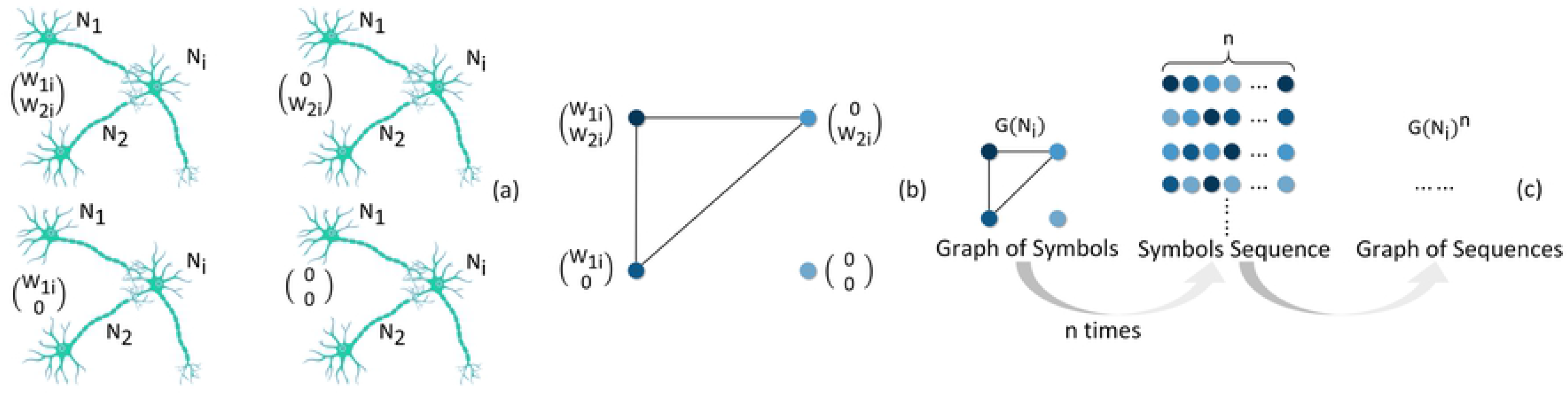
Visualizations of neural information space and graph. (**a**) We define that 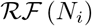 consists of 2 types of neurons with unique tuning curves. Assume that each *j* type includes only one neuron, which is marked as *N*_*j*_. We know that 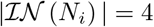 and all the possible cases are given above. (**b**) Assume that both of the 1st and 2nd type of neurons can activate *N*_*i*_ independently. Thus, the connected component of 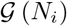 contains only the neural information cases where one or both of the 1st and 2nd type of neurons spike. As for the rest cases, they are isolated nodes. (**c**) The construction of 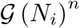 based on 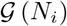.

We address two main questions about the neural information space 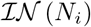: How many elements can there be in the space? What is the topology structure of the space?

For the first question, if *C* maintains constant, the variability of 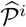 will be the determinant of the cardinal number of the space. We propose a method to work out 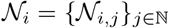, which is the set of all neural types classified based on tuning properties (the section “The cardinal number of spike-based information space” in Methods). Ideally, each type of neurons might all emit spikes or all keep silence when a given stimulus comes in. Thus, the total variation of neural spiking states is measured as 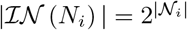 (see the section “The cardinal number of spike-based information space” in Methods. As for the potential-based information, 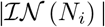 is also measured in the section “The cardinal number of potential-based information space” in Methods).

For the second question, the topology concerned here is relevant with information confusion. In Shannon’s theory, there is a confusion relation between two messages if each of them might be mistook for the other. For neural systems, we define that two neural information cases are confused with each other if and only if they can induce same neural response of the receiver neuron. Specifically, the neural response of receiver neuron is demanded explicitly to be spiking in our research. Thus, we represent the confusion relationships based on a binary tuple 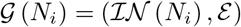, where 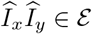 if and only if both 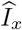 and 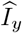 can make *N*_*i*_ spike when *N*_*i*_ is not in the refractory period (see the section “Potential-based information graph definition” in Methods). **Fig. 2a-b** show an example of 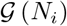. In our framework, the proposed information confusion is naturally determined by the interactions between synaptic connection states and neural tuning properties, which does not rely on noises.

#### Neural zero-error capacity

In information theory, zero-error capacity is used to indicate the supremum of information transformation rates with zero probability of error in a given channel. Following the classical definition, we define the neural zero-error capacity Θ (*N*_*i*_) as

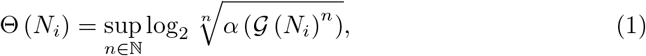

where *α* is an independence number (see the section “Zero-error capacity definition of the neural information space” in Methods). In (1), 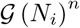 can be understood as a neural information graph corresponds to the process during which 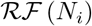 sends message to neuron *N*_*i*_ via the channel *n* times (**Fig. 2c** and the section “Zero-error capacity definition of the neural information space” in Methods). In other words, we can treat 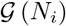 as the graph that indicates the relations between basic neural information symbols, and 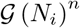 is the graph for the relations between the neural information sequences constructed by *n* symbols. Apart from that, in the neural information graph, the existence of edge between two nodes means that there is a confusion relation between them. Thus, the independence number *α* measures the maximum number of neurons can be picked from the graph and have no confusion with each other (see **Fig. 3a** and the section “Zero-error capacity definition of the neural information space” in Methods). To sum up, Θ (*N*_*i*_) measures the supremum of information rate can be sent from 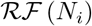 to *N*_*i*_ with no confusion.

**Fig 3.**
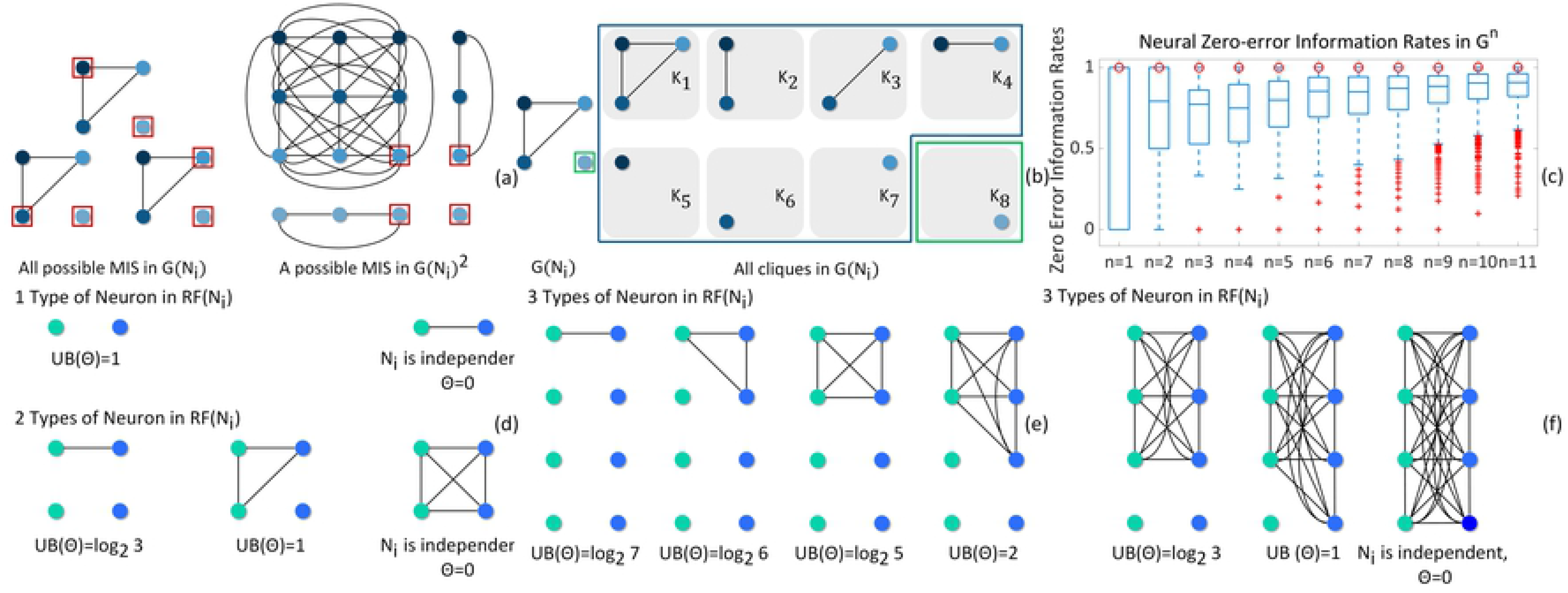
An example of the neural zero-error capacity measurement. (**a**) We mark the maximum independent set (MIS) in 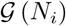 and 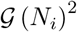 with red boxes. (**b**) It is clear that there are 8 cliques in 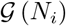 in total and all of them have been given above. As for 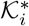, since there is an isolated node (marked by a green box) in the graph and it can be only included in one clique, which is *K*_8_, we can conclude that *μ* = 1. Similarly, *θ* = 1 since there is no other clique in this connected component. (**c**) For the 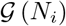 in **Fig. 2b**, we randomly search the zero-error transformation cases in 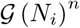 with *n* = 1, …, 11 by solving the independent set searching problem. For each 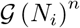, we randomly pick its independent set 1000 times to obtain a set of zero-error information rates. Moreover, based on our theory, an upper bound of Θ (*N*_*i*_) can be predicated (marked by red circles), which equals 1. (**d-e**) The upper bound is marked as “UB”. *N*_*i*_ is independent means that its spiking state is independent from spiking states of the neurons in 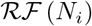, which indicates that *N*_*i*_ must be an input neuron rather than an intermediary neuron.

An important property of equation (1) is that the neural zero-error capacity of any given neural information graph has a close relation with the maximum clique assignment *λ* of the graph. In our work, we suggest that any neural information graph 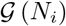 has a maximum clique assignment *λ*_*i*_ determined by the properties of 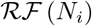 (see the section “The maximum clique of neural information graph” in Methods, (24) and (25)). Then, because of the independence number *α* satisfies *α* ≤ λ^−1^, we can tell that *λ* can offer an upper bound for the neural zero-error capacity, which is given as Θ (*N*_*i*_) ≤ log2 *λ*_i_^−1^ (see the section “Upper bound estimation method” in Methods, (22)). Thus, combine the results in (24) and (25) with (22) in the section “Upper bound estimation method” in Methods, we can know for any neuron *N*_*i*_, there is

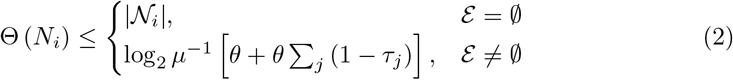

where *τ*_*j*_ indicates if 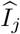 can activate neuron *N*_*i*_. *τ*_*j*_ equals 1 when 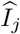 can make *N*_*i*_ spike and equals 0 when 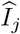 can not.

We need to pick one node from 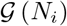 that has minimum degree (the number of cliques that include this node is smallest), let 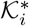 be the set of all cliques that contain this node, and let 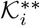 be the set of all cliques in the same connected component with this node. Then, define 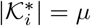 and 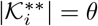. Based on (33), (34) and (35) (see the section “The maximum clique of neural information graph” in Methods), *μ* and *λ* can be easily calculated. **Fig. 3b** is an example of the calculation. More examples can be found in **S1 Fig** in Supporting information.

When 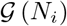 is not a complete graph, the upper bound predicted above can be proved as the supremum. As for the case when 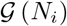 is a complete graph, Θ (*N*_*i*_) = 0 can be directly obtained, thus the upper bound measurement is not necessary (see the section “Upper bound estimation method” in Methods). Therefore, our method can measure the limitation of the zero-error transformation rate efficiently in any possible case.

In real neural systems, the situations with 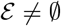 are ubiquitous. Thus, we suggest that the limited neural zero-error capacity described by (2) is of great significance in understanding the limitation of neural information processing. Shannon had proved that the zero-error information rate of a given channel cannot be decreased by lengthening the message sequence (in other words, by increasing *n* of 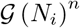 [23]. In **Fig. 3c**, we demonstrate a sample of the neural zero-error information rate measurement to show that the neural zero-error information rate is not decreasing either (see the section “Upper bound estimation method” in Methods). More examples can be seen in **S2 Fig** in Supporting information. In addition, many other basic cases of the neural information graph and its neural zero-error capacity or the corresponding upper bound are listed in **Fig. 3d-f**.

Up to now, we have introduced our neural information confusion theory based on the spike-based information. For a neuron, if the neural zero-error capacity is reached by the transformation rate, then there must exist information confusion; if it is not reached, there still exists the probability of confusion. This theory can be applied to any kind of finite and discrete neural information. To demonstrate its potential application in neuroscience studies, we extend our analysis to a simulated neural network and show how the information confusion affects the coding process.

### Information confusion in the coding process

#### Neural information confusion detection

During neural coding, information is transmitted between neurons and frequently but not necessarily involved with the information confusion. Apart from that, it is known that the neural response preference contained in the neural information is summed and transmitted as well, which usually implies a selectivity generalization along the neural pathway (e.g., the selectivity generalization from the V1 neurons to MT neurons in the visual system [36–38]). Due to the similarity between information confusion and selectivity generalization, it might be misunderstood that they are two equivalent conceptions. Here, we propose a method to directly detect the information confusion in neural coding and distinguish it from the selectivity generalization.

We demonstrate a dyeing experiment involves with a network of 500 neurons. In the experiment, the network is assumed to code a stimulus sequence consists of different vectors. Each type of input neuron with unique stimulus selectivity is marked with a specific color and all intermediary neurons are initialized with no innate preference and marked as white (**Fig. 4a**). For simplification, we use numbers to index those vectors in our results (**Fig. 4c**). In each iteration, the intermediary neurons are dyed depending on the relation between the neural responses of it and its pre-synaptic neurons (see **Fig. 4b** and the section “Dyeing method for visualizing the existence of selectivity generalization phenomenon” in Methods). Thus, we can see how the selectivity is generalized through the information transformation process (see **Fig. 4d**. And more examples can be seen in **S3a-d Fig** in Supporting information). The corresponding spiking states of this neural population can be seen in **Fig. 4e**, where we compare the observed responses of intermediary neurons with the tuning curves of 2 types of input neurons. We show that many intermediary neurons show responses to the stimuli out of the stimulus range preferred by any single type of input neurons, which offers verification for the results in **Fig. 4d**.

**Fig 4.**
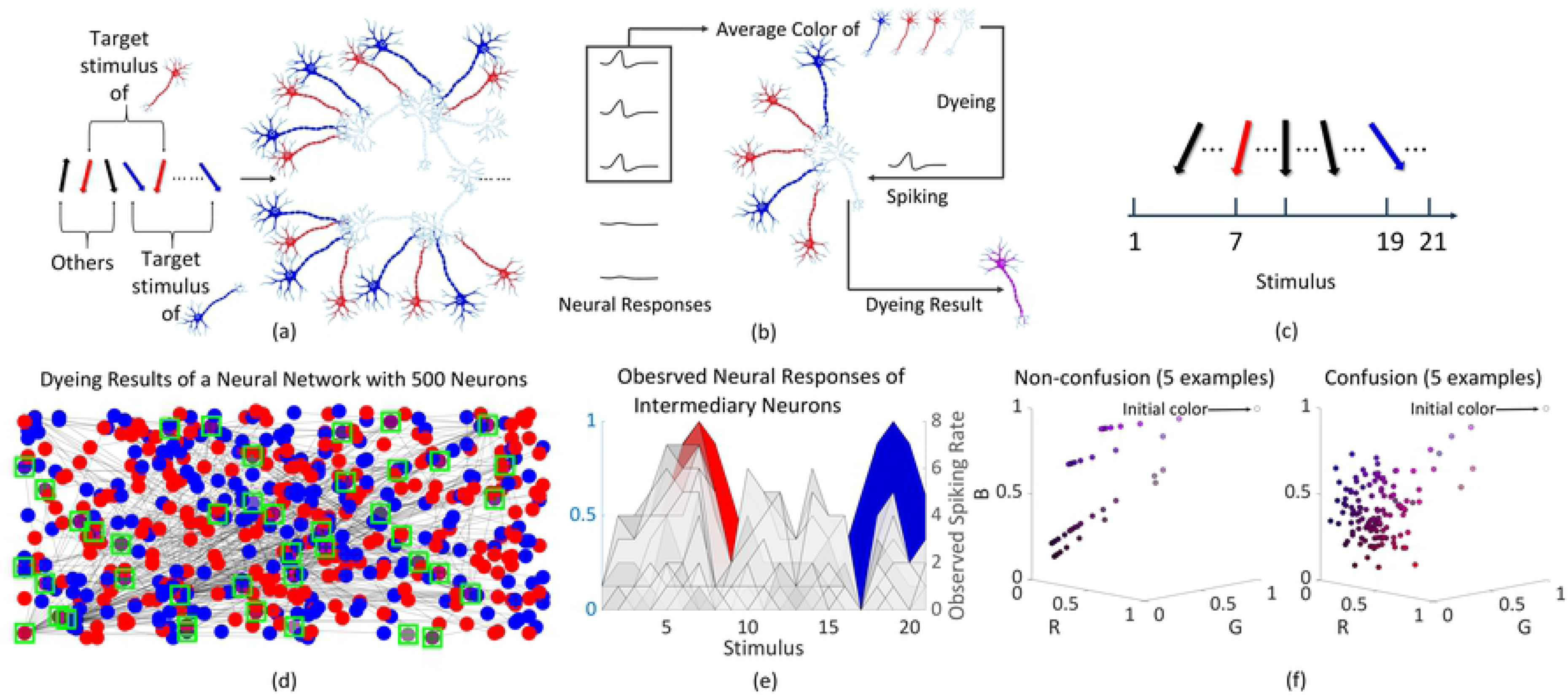
Detecting neural information confusion during neural coding. (**a**) There are 2 types of input neurons (respectively marked as pure blue and red). The stimulus sequence consists of various vectors, among which, the blue and red vectors are respectively preferred by the input neurons with same color. (**b**) In each iteration, if an intermediary neuron spikes, then it is dyed with color averaged from its previous color, and the colors of the lately spiked neurons in its receptive field. Thus, if the color of an intermediary neuron is bulish-red (the color has non-zero red and blue components), then it has multi-stimulus preference. (**c**) The index of stimulus. (**d**) The dyeing result after coding the given stimulus sequence in 100 iterations. (**e**) The red and blue areas respectively stand for the normalized tuning curve of the input neurons with same color. The gray areas measure the response of intermediary neurons. (**f**) The information confusion can be detected based on the color variation trajectory.

We know that the existence of neural information confusion means that different neural information implies same neural response of the receiver neuron. In the dyeing experiment, each spiking intermediary neuron is dyed with the color averaged from its previous color and the colors of the lately spiked neurons in its receptive field. So, if there is only one message that makes this neuron spikes, then its color will straightly approach to the averaged color of the spiking neurons described by this information (with a straight trajectory). Otherwise, if the variation trajectory of its color is winding or oscillating (not only one message can activate this neuron), then we suggest that there exists the neural information confusion (see the section “Detect the neural information confusion based on the dyeing experiment result” in Methods). We use the experiment data to show several examples for the color variation trajectory with/without confusion in **Fig. 4f**. More examples can be seen in **S3e-f Fig** in Supporting information. As for the selectivity generalization, it only requires the color has at least two non-zero color components (**Fig. 4b**) and the variation trajectory can be either straight or winding. Together, we conclude that the information confusion is a special case of the selectivity generalization with a winding color variation trajectory and thus they are not equivalent conceptions.

#### The effects of information confusion

After detecting the existence of information confusion during the coding process, a natural thought is to wonder if the confusion affects neural coding. To answer this question quantitatively, it is necessary to review three main parameters widely used in neural coding studies, which are the total response entropy *H* (measures the total variation of neural response), the noise entropy *H*\* (measures the variation of neural response that cannot be explained by the stimulus) and the mutual information *H*\*\* (measures the variation of neural response that can be explained by the stimulus). They satisfy *H* = *H*\* + *H*\*\* (see the section “Calculate *H*, *H*\* and *H*\*\*” in Methods for detail). Based on the noise entropy *H*\*, we can further define 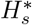 for each input stimulus *s* (measures the variation of neural response that can not be explained by *s*). The smaller 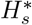 is, the less noise the coding process of *s* produces. In other words, the neural activities can be better explained by *s*. Thus, we define the stimulus subset that can better explain the neural activities as 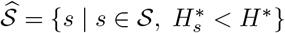. Then, we define the coding scope as 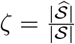 (see the section “Definition of the coding scope” in Methods) to measure the proportion of the stimulus that can better explain neural activities in all stimuli.

What interests us is if there exists information confusion, then what will happen to *H*, *H*\*, *H*\*\* and *ζ*? We demonstrate the changes of these four parameters in an experiment with a random stimulus sequence 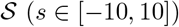. The experiment is carried out 3000 times. For neuron *N*_*i*_, we assume its receptive field 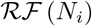 contains *k* neurons (*k* ∈ [40, 60]). In each iteration, every neuron in 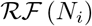 is set to has a tuning curve 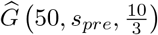, where *s*_*pre*_ ∈ [−10, 10]. And the synaptic connection between *N*_*i*_ and 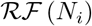 has a connection strength randomly picked from [−1, 1]. Based on those settings, we can simulate almost all the cases of neurons’ behavior in neural systems.

In the simulation, there are 357 iterations without confusion and 2643 iterations with confusion (about 88.1% iterations). When information confusion exists, we find that the neural response variability (*H*), stimulus-irrelevant neural activity variability (*H*\*) and stimulus-triggered neural response variability (*H*\*\*) are reduced when the information is transmitted from pre-synaptic neurons to the post-synaptic neurons in most cases. The average reductions are 2.871 bits (*H*), 1.963 bits (*H*\*), and 0.909 bits (*H*\*\*), and the average reduction proportions are 68.4% (*H*), 71.6% (*H*\*), and 62.3% (*H*\*\*) respectively. In contrast, the coding scope (*ζ*) usually increases after the transformation, the average increase is 0.257 bits and the increase proportions is 62%. See **Fig. 5a-b** for these results in detail.

**Fig 5.**
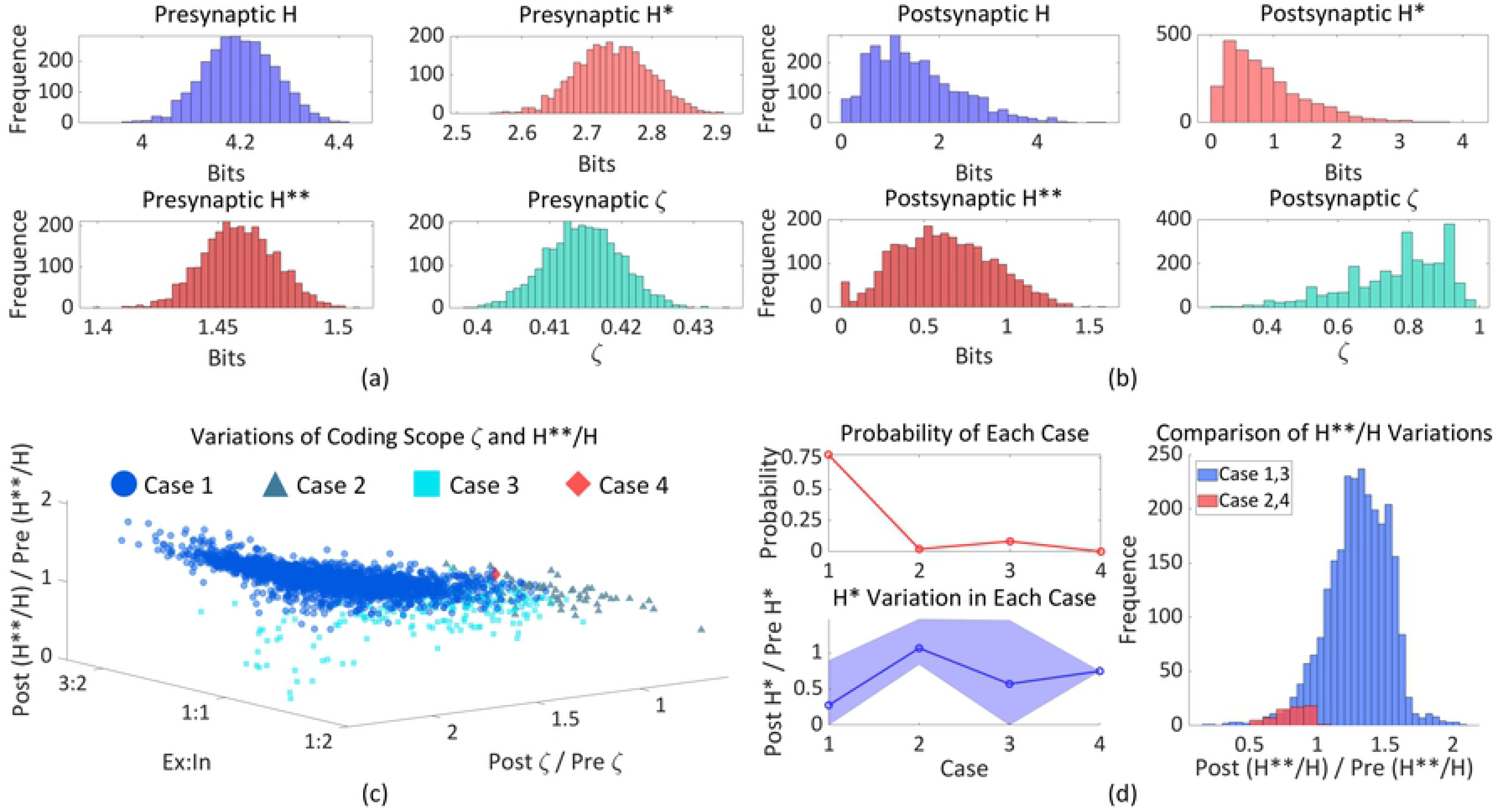
Effects of neural information confusion on neural coding. (**a-b**) The pre-synaptic frequency distributions of *H*, *H*\*, *H*\*\* and *ζ* (averaged between all pre-synaptic neurons) and the corresponding post-synaptic frequency distributions. (**c**) The variation distribution of *ζ* and 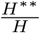. (**d**) We show the occurrence probability of each variation case and distinguish them based on the variation trend of *H*\*. Then we show the variations of 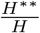 with respect to different cases.

To explore how the stimulus-related information evolve in the neural system, we use 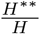 (a parameter that measures the proportion of explainable part in the total neural response variation) in the following analysis and make a direct comparison between the 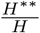 with the coding scope. Their relationship can be defined as 4 cases: **case 1** means *ζ* and 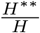 increase; **case 2** means both *ζ* and 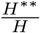 decrease; **case 3** means that *ζ* increases while 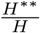 decreases; **case 4** means that *ζ* decreases while 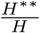 increases. In **Fig. 5c**, we show the distribution of these 4 cases with the synaptic connection state (the ratio of all inhibitory to all excitatory synaptic strength absolute values), which suggests that the synaptic connection situation cannot offer a clear discrimination between them. Thus, the variations of *ζ* and 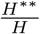 (respectively measured by the ratio of post-synaptic *ζ* to pre-synaptic *ζ* and the ratio of post-synaptic 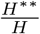 to pre-synaptic 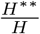) can not be simply explained by synapse-relevant factors. In **Fig. 5d**, the occurrence probability of each case is measured. It can be seen that **case 1** is the most frequently occurred case (probability is 0.778) while the other three cases are scarce. Then, we compare the variation of *H*\* (measured by the ratio of post-synaptic *H*\* to pre-synaptic *H*\*), and turns out that the cases where *ζ* increases (**case 1** and **3**) correspond to the most significant reduction in *H*\* (the ratio of post-synaptic *H*\* to pre-synaptic *H*\* is less than 3 : 5), while the cases where *ζ* decreases (**case 2** and **4**) correspond relatively slight reduction or even increase in *H*\*. Moreover, we also compare the variation distribution of 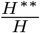 between the cases with coding scope increase and decrease, showing that 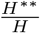 has accordant variation trend with the coding scope *ζ* (increase or decrease together). Another experiment result can be seen in **S4 Fig** in Supporting information.

To sum up, the information confusion has both detrimental and beneficial effects on neural coding. On the one hand, the total variability *H* of neural responses is reduced, which limits the mutual information *H*\*\*; On the other hand, it is highly possible that *H*\* decreases when the stimulus-irrelevant neural activities are controlled and both *ζ* and 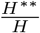 increase, which means that the interpretability of neural responses is improved and the activities can be better explained by a wider range of stimuli.

## Discussion

### Summary of our work

In the present study, we reveal an innate limit of the information processing in neural system by indicating the pervasive information confusion phenomenon during the information transformation between neurons.

We first define the neural information confusion in the content of neuroscience and propose a practical method to work out the upper bound of the information transformation rate with zero error of any given neuron (neural zero-error capacity). This systematic theory can be applied to any kind of discrete and finite neural information (e.g., spike-based or potential-based). For a neuron, if the neural zero-error capacity is reached, then there must exist neural information confusion; if it is not reached, there still exists the possibility of neural information confusion.

We then propose a practical method to analyze the effects of information confusion on neural coding. The results suggest that the effects of information confusion can be either detrimental or beneficial. On the one hand, it controls the total variability *H* of neural responses and limits the mutual information *H*\*\*. On the other hand, it improves the interpretability of neural responses as that the stimulus-irrelevant neural activities are controlled.

As unveiled in our work, the precision limit caused by information confusion features widespread presence during the neural information processing, and is intrinsically determined by neural tuning properties and synapse states. This innate limit plays a critical role in characterizing the neural coding process, leading to the variation of coded information along the information transformation pathway. In sum, we demonstrate that neural system is not an inherently error-free information processing system even under ideal conditions, and its essential limit in information transformation creates significant effects in neural coding.

### Neural information confusion and relevant topics

Besides information confusion, there are three other resources related with the information limitation of neural systems. The first one is the information loss caused by channel capacity [13–18]. Information loss happens when the entropy of information exceeds the channel capacity (the maximum transfer rate that the channel supports). Information loss concerns about the limited information transfer rate rather than the transfer precision. So it is more related to the reduction of the total response entropy *H* in our research [19–22].

The second one is the information limiting correlation observed experimentally. The origin of it still remains controversial. A classical perspective argues that the similarity in tuning properties implies the correlations among neurons for a target stimulus and affect neural coding significantly [39–41]. These positive correlations are inevitably caused by the shared input connections between neurons with similar tuning characteristics, which suggests that shared connections between neurons might lead to limited information [42]. Recent studies have contradicted this hypothesis in several aspects. Rather than originate from the shared connections, the limitations are suggested to be caused by the correlations proportional to the product of the derivatives of the tuning curves. They spontaneously emerge in the finite information encoding and storage process of a sufficiently large neural population [43–45]. Despite the controversy on the origin mechanism, it is confirmed that the correlation can limit information in neural coding (similar with the information loss). A similar finding is proposed in our research, which suggests that compared with the pre-synaptic variability (information) of neural responses, the post-synaptic one is frequently controlled.

The third one is the information error (e.g., errors caused by the channel unreliability [19] or random noises [20–22]). Compare with information confusion, there are two main differences. The first difference lies in that the information confusion does not focus on the noises added to the neuron, confusion is determined congenitally by the tuning properties, even in an ideal situation with no external noise. The second difference is that the transfer losses and errors happen during the transformation while the confusion happens after the information arrives at the receiver.

## Methods

### Leaky integrate-and-fire networks

Here we describe the network with leaky integrate-and-fire neurons demonstrated in our research.

#### Network definitions

In our paper, the recurrent network of leaky integrate-and-fire neurons is used to create the electrodynamics involved in the neural information transformation process. Rather than actuate all neurons directly based on the stimulus as previous researches did [43], we distinguish input neurons from the neuron set and define the stimulus as the synaptic drive for those neurons.

The basic element in our simulation is the differential equation for the membrane potential time evolution of leaky integrate-and-fire neurons with current synapses, which is defined as

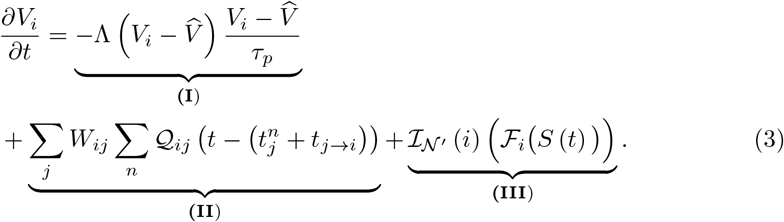

In this equation, item **(I**) is the leak current. Λ denotes the Heaviside step function, *τ*_*p*_ is the leaky membrane time constant and 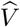 is the resting potential. If the membrane potential *V*_*i*_ is not less than 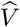, then neuron *N*_*i*_ will be involved in a hyperpolarization process described based on the leaky mechanism.

**(II**) is the recurrent item, which consists of the spiking mechanism and the synaptic input. In this item, *W* is the weighted adjacent matrix that defines the connection strength between neurons,

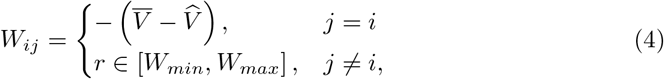

where *W*_*min*_ and *W*_*max*_ are respectively the minimum and maximum connection strength, and 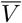 is the spiking threshold. And 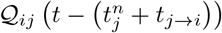 is the synaptic response to the spike, which is given as

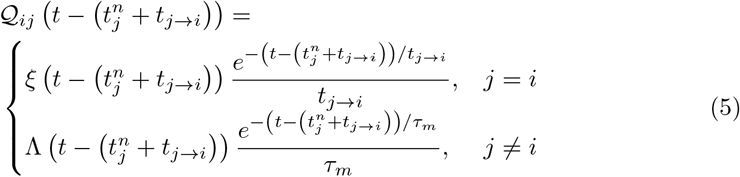

in which *τ*_*m*_ is the membrane time constant. And *ξ* is given as

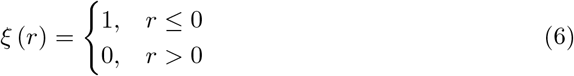

In equation (5), 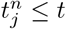 denotes the time of the *n*th existing spike of neuron *j*. And *t*_*j*→*i*_ measures the time cost, if *j* = *i*, then *t*_*j*→*i*_ is the refractory period; If *j* ≠ *i*, then *t*_*j*→*i*_ is the average transmission delay of the spike. Based on equations (4) and (6), 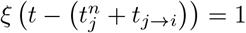 if and only if 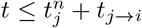, which means the neuron is still in the refractory period. Furthermore, based on equations (4) and (5), it is clear that if *j* ≠ *i*, then 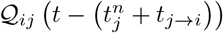 corresponds to a current pulse with the random connection strength *r* selected based on an uniform distribution. As for the case with *j* = *i*, if the membrane potential of neuron *i* reaches the spiking threshold 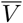, then it emits a spike, after which it experiences a polarization and the voltage is reset to 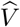.

**(III**) is the stimulus drive item, whose definition is learned from [43]. In this item, 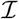 is the indicative function and 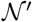 is the index set of all input neurons.

If neuron *i* is an input neuron, then 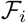 is the synaptic drive of it based on the stimulus *S*. The definition of 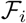 can vary based on research targets. In our research, the synaptic drive is defined to represent the characterized response preference of input neurons. More specifically, each input neuron *N*_*i*_ has its own tuning curve *G*_*i*_, which decides its response strength to given stimuli (see the **subsection** in Methods for details). Then, we let 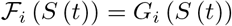, thus the stimulus preference of each input neuron is defined.

#### Network parameters

Here we present the simulation parameters for the recurrent network. Note that none of the results depend critically on the parameter values. Generally any bio-plausible setting for the neuron populations can be considered.

**Table 1.** Parameter settings for networks

| Par | Value | Par | Value |
| --- | --- | --- | --- |
| $\widehat{V}$ | 0 | $\overline{V}$ | 5 |
| $W_{min}$ | -0.5 | $W_{max}$ | 1 |
| $\tau_p$ | 20 ms | $\tau_m$ | 2 ms |
| $t_{j \rightarrow i}$ | 3 ms | $t_{i \rightarrow i}$ | 2 ms |

### Neural information confusion and neural zero-error capacity

In this section, we propose a practical method to measure the upper bound of neural zero-error capacity directly based on the properties of the given neuron.

#### Tuning curve and neural response

In our research, the stimulus 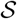 is set as a sequence where each 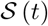 is selected randomly and uniformly from the stimulus interval 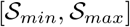.

The tuning curve of any given input neuron is defined as 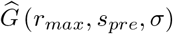, in which the preferred stimulus *s*_*pre*_ is randomly selected from 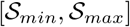 based on a random distribution *F*. In our research, *F* is set to be an uniform distribution for simplification. And the maximum response rate *r*_*max*_ is randomly selected from an empirical interval [40, 60] based on an uniform distribution. *σ* represents the width of the tuning curve, which is set randomly from 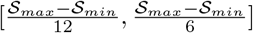.

In detail, the mathematical definition of the tuning curve is given as

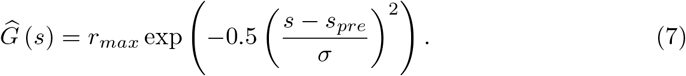

As for the intermediary neuron, its estimated tuning curve can be calculated based on the recurrent connection strength matrix *W* and the tuning curves of input neurons. Assuming that each input neuron *N*_*i*_ has a tuning curve 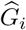, the tuning curve 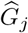 of a given intermediary neuron *N*_*j*_ is estimated as

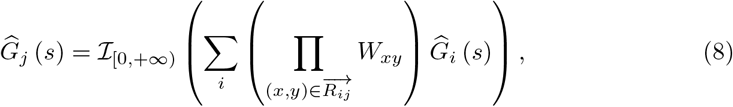

where 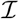 is the indicative function and 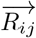 is the shortest route from *N*_*i*_ to *N*_*j*_. Based on equation (8), we actually let the tuning curve of intermediary neuron be shaped by the postsynaptic potentials of input neurons.

#### Neural information space definition

There are two kinds of widely used neural electrical information forms. The first one is the spike-based and the second one is the potential-based. The spike-based information has 2 symbols (spiking and non-spiking) while the potential-based information has multi-symbol (the number of symbols is determined by the number of all possible potential states that a neuron can have). For simplification, we use the spike-based information to introduce our theory, yet all the necessary definitions for both types of information will be given as following.

##### Spike-based information space

To define the spike-based information space, it is necessary to define the neural spiking states in the receptive field of each given neuron. For a neural population with *n* neurons, we define that each neuron *N*_*i*_ is equipped with a spiking state 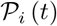 at moment *t*, in which

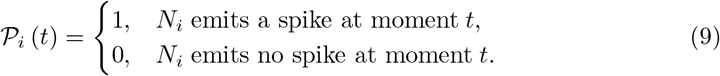

Then, the spiking states vector 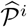 within the intra-system receptive field of any neuron *N*_*i*_ is given as

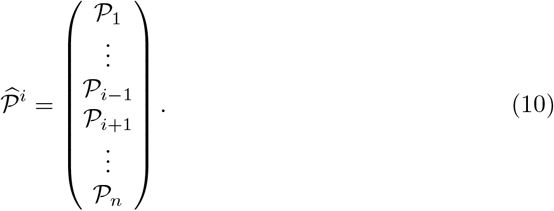

In equation (13), we exclude the spike state of *N*_*i*_ itself to leave out the recurrent information. Analogously, we can also define a non-recurrent connection strength matrix *C* to indicate the synaptic connections between neurons by exclude the recurrent items in *W*. More specifically, we leave out the all elements on the primary diagonal of *W*

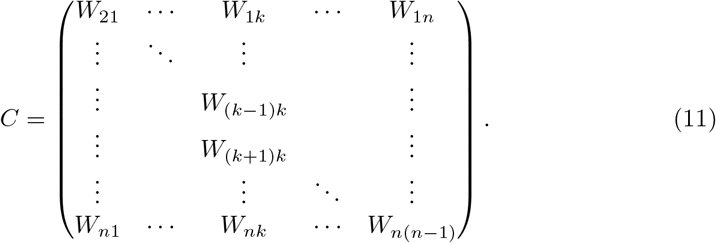

Let *C*_*i*_ be the *i* column of *C*, by calculating the Hadamard product ⊙ of 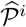 and *C*, we can represent any possible case of the spike information that *N*_*i*_ can receive, based on which, the spike-based information space 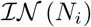 is defined as

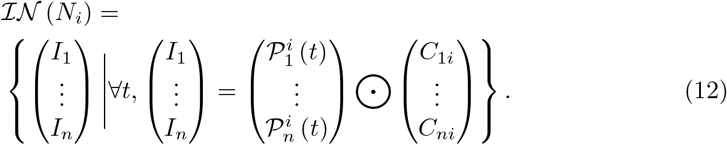

##### Potential-based information space

The main difference between spike-based and potential-based information lies in that the first one only indicates whether a neuron spikes or not while the second one can not only show the spiking state, but also indicate the membrane potential.

For a given neuron *N*_*i*_, based on (3), its potential state is *V*_*i*_ (*t*). When the potential state reaches to 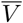, there will be a spike. In real experiment, the measurement technology can not realize arbitrary precision, there must be a precision limitation of it. So, we can treat *V*_*i*_ (*t*) as discrete.

Following the idea used in the definition of the spike-based information space, we define the potential states vector 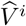 within the intra-system receptive field of any neuron *N*_*i*_ is given as

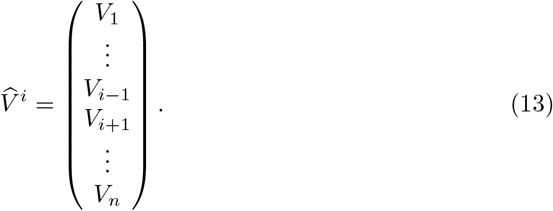

And the corresponding potential-based information space is given as

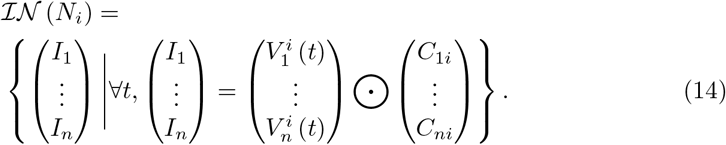

#### Measure the number of elements in the neural information space

##### The cardinal number of spike-based information space

After defining the spike-based information space 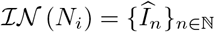, a natural thought that comes to mind is how many elements there might exist in it. Since similar tuning properties usually result in the similar responses to given stimuli, we can reasonably measure the total variation of neural spiking states based on the characterized response preference. More specifically, we use the Wasserstein distance to indicate the differentiation between the tuning curves of two neurons *N*_*x*_, *N*_*y*_, which is given as

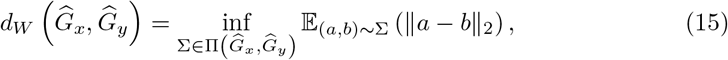

where 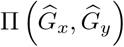 is the set of all possible joint distributions of 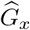 and 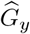. For each given joint distribution Σ, equation (15) takes one sample (*a, b*) from it at each time and eventually works out the expectation of *L*_2_ norm || · ||_2_ of all samples. Then, we use the infimum of all possible expectation values to represent the distance between 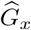 and 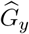.

Based on this definition, we define an equivalence relation ~*w* that *N*_*x*_ ~*w N*_*y*_ if and only if 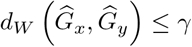, in which *γ* is a given threshold. Thus, for neuron *N*_*i*_, its intra-system receptive field 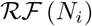 can be classified into 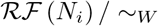, where each element is a neuron type with specific characterized response preference. In our research, we mark that 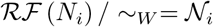, where 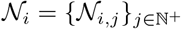. If we let *γ* approach to 0, we know that the homogeneity between same type neurons will be increased. Thus, under ideal conditions, we assume that for each type of neurons, they can either all emit spikes or all keep silence when a given stimulus comes in. As a result, the total variation of neural spiking states in the intra-system receptive field of neuron *N*_*i*_ is defined as

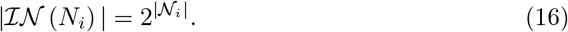

##### The cardinal number of potential-based information space

Following the idea discussed previously, we continue to use the equivalence relation ~*w* to obtain 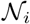. We assume that the measurement accuracy limitation of the membrane potential is Δ*V*. Thus, for any given neuron *N*_*i*_, the number of all possible potential states of it can be measured as 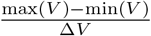, where max (*V*) is the maximum membrane potential and min (*V*) is the minimum one. So, the total variation of neural spiking states in the intra-system receptive field of neuron *N*_*i*_ is defined as

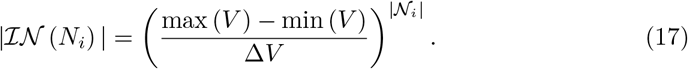

#### Neural information graph definition

##### Spike-based information graph definition

Up to now, we have defined the spike-based information space and analysed its cardinal number. It is time to turn to the topology structure of the space. In our research, the concerned topological relation is related to confusion. For nervous system, we define that two spike-based information cases 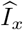 and 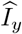 of 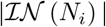 are confused with each other if and only if both of them can make *N*_*i*_ spike when *N*_*i*_ is not in the refractory period. To be more specific, we define an equivalence relation Δ*C* to represent the confusion relation mentioned above, which is given as 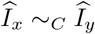 if and only if

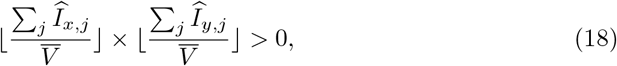

it is clear that (18) means that both of the postsynaptic potentials correspond to 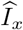 and 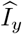 can reach to the spiking threshold 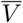 of *V*_*i*_.

To seek for a better representation of the confusion relation, we propose the definition of spike-based information graph. For each given neuron *N*_*i*_, its spike-based information graph is defined as

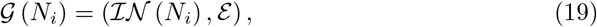

where 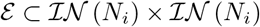, and 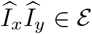 if and only if 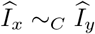.

Here we suggest that each connected component in 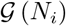 must be a clique, or in other words, a complete subgraph. The proof is trivial since ~*C* is an equivalence relation and has transitivity.

##### Potential-based information graph definition

The potential-based information graph is very similar to the spike-based one. The only one difference between them is about the definition of the equivalence relation ~_*C*_.

At first, we define a equivalence threshold *ε*_*e*_ > Δ*V*. For two potential-based information cases 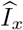 and 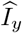, let us assume that the membrane potential states of *N*_*i*_ receive them are 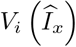 and 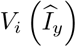, both of which can be calculated based on (3).

Then, if

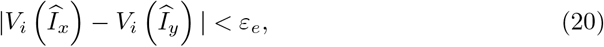

then we treat the membrane potential states described by 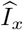 and 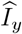 are same as each other.

Then, we can define that

- if 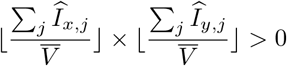, then 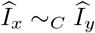;
- given 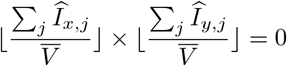, if 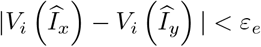, then 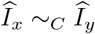.

Based on the above definition, we can follow the method in (19) to define the potential-based information graph.

#### Zero-error capacity definition of the neural information space

Up to now, we have obtained the neural information space and the corresponding graph (both the spike-based and the potential-based). We then turn to measure the zero-error capacity based on the graph. Note that what we will discuss is independent from the selection of neural information type, so we do not distinguish between different types of neural information.

In the information theory, if we treat 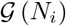 as the representation of relation between symbols (nodes in the graph), then 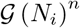 represents the relation between the sentences consists of *n* symbols. We know that if there is an edge between two nodes, then the symbols or sentences represented by them are easy to be confused with each other. For informatics, confusion is a kind of error. So, a meaningful question is to ask about how many symbols or sentences from a given system can be transmitted with zero-error at most. To answer this question, Shannon defined zero-error capacity [23], which is given as

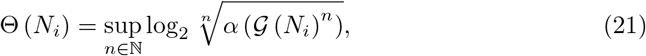

where *α* is the independence number, which indicates how many nodes can be included in the maximum independent set of 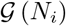.

In this section, we will not go deeper into the discussion of how Shannon create this concept step by step. What we want to emphasize is that Shannon and later researchers [23, 26, 27], discovered that for any graph 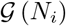

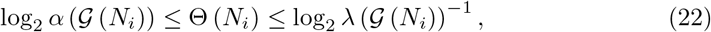

where 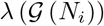 is the maximum clique assignment. Based on this inequality, when it is hard to work out Θ (*N*_*i*_) directly, we can still obtain a bounded measurement of it.

#### The maximum clique of neural information graph

A relevant concept of the neural information graph is the maximum clique assignment. In graph theory, the maximum clique assignment of a graph *G* is defined as

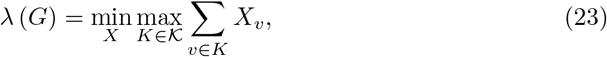

where *X* is any random distribution *X* = {*X*_*v*_ | *v* ∈ *V*} and 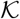 is the set of all cliques in the graph.

In our research, the maximum clique assignment *λ*_*i*_ of any given 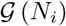 satisfies

- if 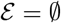, then

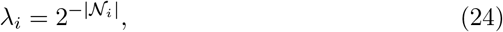
- if 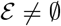, then there is

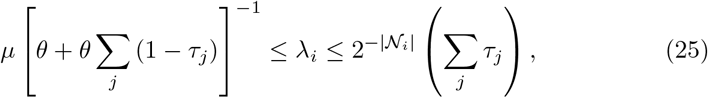

where we pick one node from 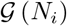 that has minimum degree (which also means that the number of cliques include this node is smallest). We then let 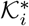 be the set of all cliques that contain this node, 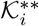 be the set of all cliques in the same connected component with this node. Next, define 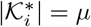 and 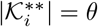. Apart from that, 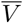 is the spiking threshold and 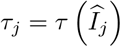 is used to indicate whether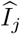 can activate neuron *N*_*i*_, which is defined as

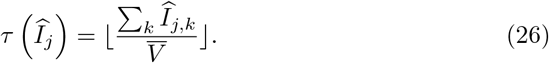

It is clear that 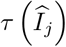 equals 1 when 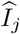 can make *N*_*i*_ spike and equals 0 when 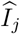 can not.

To provide a better understanding for (24) and (25), we give the following proof for this theorem.

##### Proof 1

*For* 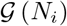, *we know*

- *if* 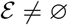, *we can know that* ∑ _*j*_ *τ*_*j*_ > 1. Then, based on equation (23), *we assign* 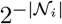 *to the nodes in* 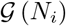, *and assign* 0 *to the edges in* 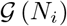, *meaning that*

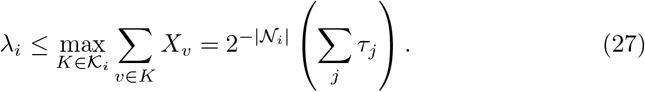 *Then, we can also consider the dual problem of the definition of λ, which is given by*

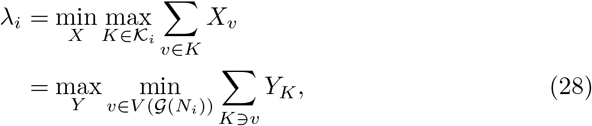

*where Y is a any random distribution* 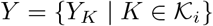. *The assignment of Y is a little tricky, which is given as*

– *first, we assign 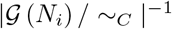 to each connected component in* 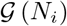. *For instance, if there are 4 connected component in* 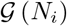, *each of them is assigned as 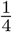. Moreover, since* 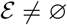, *we know that the number of connected components can be worked out by* 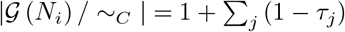 ;
– *second, for a connected component containing k cliques, assign each clique as* 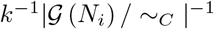. *Based on the assignment described above, it is easy to know*

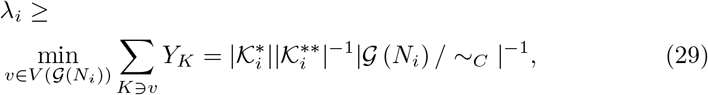

*which can be rewritten as*

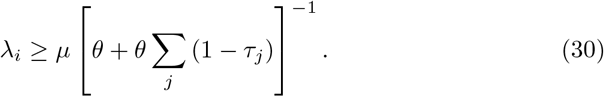 *Thus, based on equation (27) and (30), we can prove that (25) is right when* 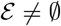.
- *If* 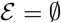, *we know that* ∑ _*j*_ *τ*_*j*_ ≤ 1 *and there is no edge in* 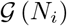. *So, the maximum clique exists in this graph is each individual node itself. Then, following the second assignment method we use above, we can also prove*

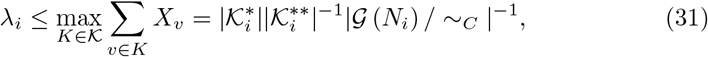

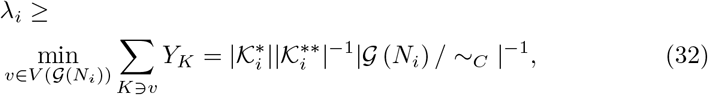

*where* 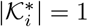, 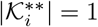 *and* 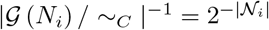. *Thus, (24) is correct when* 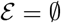. *Here we don’t repeat the proof in detail*.

Specially, to make the calculation easier, for (30), we can have following analyses for *μ*

- 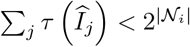 means that not all neural information cases can make *N*_*i*_ spike and 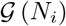 is not a complete graph. Under this condition, we know there must be at least one isolated node in the graph, and it implies that

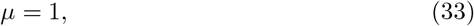

since the isolated node is included in only one clique, which is the node itself.
- 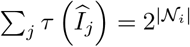 means that any neural information case can activate *N*_*i*_, so 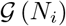 is a complete graph. Under this condition, every node is included by same amount of cliques. For convenience, we mark 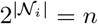. We then randomly pick one node from 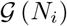, as for its corresponding 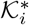, we know that

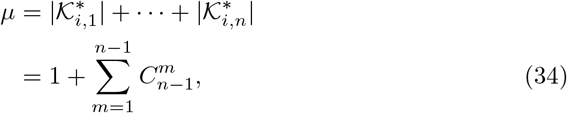

where 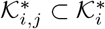 is the set of all the cliques that include this node and have *j* nodes in total.

We can also tell that the *θ* in equation (30) can be worked out by

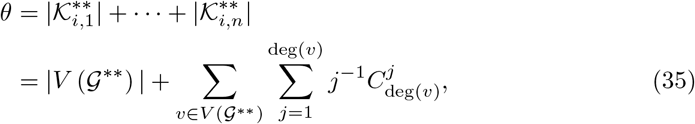

where 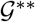 is the given connected component and deg (*v*) measures the degree of *v*. An important property is that if *μ* = 1, which means that the connected component is a isolated node, we can know *θ* = 1 as well since there is only one clique in the connected component (the node itself).

#### Upper bound estimation method

Based on (24) and (25), we can simply deduce that for any given 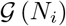

- if 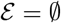, then

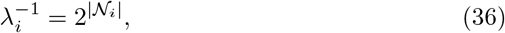
- if 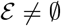, then

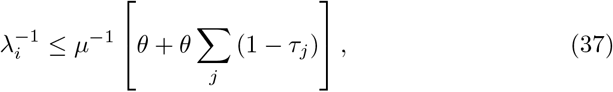

so combine (36), (37) with (22), we can know

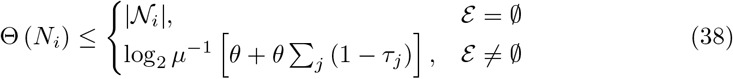

An important thing is that (38) is actually the supremum of the neural zero-error capacity if 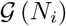 is not a complete graph. The following is the proof

#### Proof 2

*If* 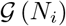 *is not a complete graph, then we know* 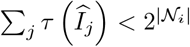, *which can be divided into two cases*

- 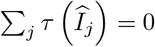, *which means that* 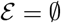 and there is no edge in the graph. Under this condition, the biggest independent set in 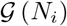 *is itself, which deduces that* 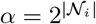. *So, based on (36), we can know that*

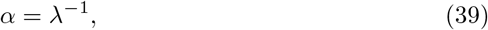

*combined with (22), (39) implies that*

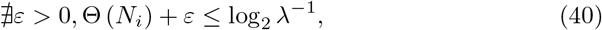

*thus the upper bound predicted by us is the supremum when* 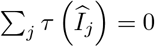.
- 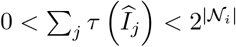, *which means that* 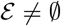 *and there must be at least one isolated node in the graph. It is clear that the isolated node has the smallest degree, being* 0. *Based on (33), we know that μ* = 1 *under this condition. Similarly, θ* = 1 *can be obtained by (35). Then, (37) can be written as*

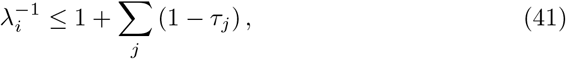

*where we know that* ∑ _*j*_ (1 − *τ*_*j*_) *measures the number of isolated nodes. And each isolated node is a connected component. As for the un-isolated nodes, they are connected with each other and belongs to one connected component. Thus, the number of connected component number of* 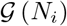 *is* 1 + ∑*j* (1 − *τ*_*j*_). *Since the maximum independent set contains only one node from each connected component, we can know α* = *λ*^−1^. *Thus, by following (40), we can prove that the upper bound is actually the supremum when* 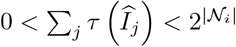.

*To conclude it, we know that when 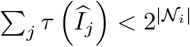, the upper bound is the supremum*.

As for the case when 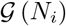 is a complete graph, we can directly know that Θ (*N*_*i*_) = 0 since you can never pick two nodes that have no edge between them in the graph. So, the measurement of upper bound is no longer needed.

#### Neural zero-error capacity definition for the dynamical transformation process

In our paper, we suggest that the synaptic plasticity mechanisms can also be included in the definition of neural information space to realize the dynamical information transformation process with plasticity.

Let us consider a plastic neural channel that transmits information to neuron *N*_*i*_ in a long enough interval [*t*_1_, *t*_*n*_]. The neural plasticity adjusts the synaptic connection strength described by *C*_*i*_ dynamically, thus the neural information space 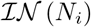 and all the concepts dependent on it change through the interval. On the one hand, at each moment, we can find a specific state for all those issues. Therefore, this dynamical information transformation process can be described by a sequence of states 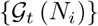, where each 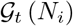 corresponds to the state at moment *t*. At each moment *t*, 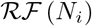 sends a symbol in 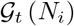 to *N*_*i*_. On the other hand, the whole process can also be treated as that 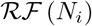 sends a *n*-symbol-message in 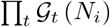 to *N*_*i*_. To summarize, while the channel state might be changed by plasticity over time, at each moment we can still obtain a corresponding static state. So, we can use a group of static neural information graphs to construct a dynamical neural information transformation process.

Following the second perspective, the neural zero-error capacity can be defined dynamically by replacing 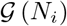 as 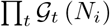 in (1), and its upper bound is given as

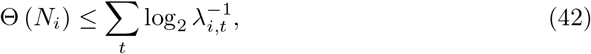

where each *λ*_*i,t*_ can be measured following (24) or (25).

### The effect of neural information confusion on the coding process

This part is concerned with the experiment methodology related to the coding scope narrowing phenomenon in our study.

#### Dyeing method for visualizing the existence of selectivity generalization phenomenon

To offer a visualization for the selectivity generalization phenomenon, here we propose a simple method:

- First, for a neural population described by the leaky integrate-and-fire network, we assume that there exist *k* types of input neurons, each type has its unique stimulus preference;
- Second, for each type of input neurons, we use a unique color to dye them as the initial color. As for the intermediary neuron, they are all initialized as white;
- Third, in each iteration of the experiment (where the neural population is set to code a stimulus sequence), if an intermediary neuron spikes, then it is dyed with the color averaged from its previous color and the colors of the lately spiked neurons in its receptive field.

Based on this setting, if the final color of an intermediary neuron contains the color components of more than one type of neurons in its receptive field, then its color must be averaged from more than one type of spiking neurons. In other words, this neuron has acquired multi-stimulus preference and the selectivity has been generalized during the information transformation.

#### Detect the neural information confusion based on the dyeing experiment result

To offer a detection for the neural information confusion in real experiments, we propose a method that can be applied directly to the results of the dyeing experiment.

Given that in the dyeing experiment, if an intermediary neuron spikes, then it is dyed with the color averaged from its previous color and the colors of the lately spiked neurons in its receptive field. So, it is easy to know that for any intermediary neuron *N*_*i*_

- if 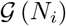 contains no edge (there is no confusion), then there are two possible cases:

– *N*_*i*_ never spikes, so the color of it always reminds to be the initial color. The variation trajectory of the color of *N*_*i*_ in the color space is a point;
– there is only one message that can activate *N*_*i*_, so the color of *N*_*i*_ gradually approaches to the averaged color of the lately spiked neurons described by this message. The variation trajectory of the color of *N*_*i*_ in the color space is straight.
- if 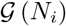 contains at least one edge (there is confusion), then the color of *N*_*i*_ will approach to the averaged color of the lately spiked neurons described by different messages in different iterations. So, the variation trajectory of the color of *N*_*i*_ in the color space is winding.

Based on those discussions, the detection of the neural information confusion can be realized by verifying whether the color variation trajectory is winding.

#### Calculate *H*,*H*\* and *H*\*\*

In neuroscience, there are three parameters relevant with the coding efficiency measurement. They are the total response entropy *H* (measures the total variation of neural responses), the noise entropy *H*\* (measures the variation of neural responses that can not be explained by stimulus) and the mutual information *H*\*\* (measures the variation of neural responses that can be explained by stimulus). In [6], researchers have proposed a practical method to calculate them, which are also used in our paper.

The first step in this method is to obtain the conditional probability distribution *P* (*r | s*) of each neuron, which denotes the probability of that the spiking rate of the neuron is *r* when the stimulus input is *s*. Of course, this distribution can be obtained directly in a real neural coding experiment. As for the computational experiment used in our paper, this distribution can be worked out based on the tuning curve, which is given as

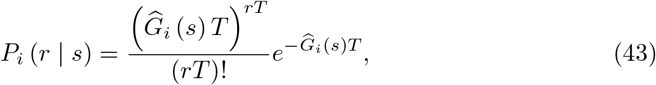

where 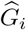 is the tuning curve of *N*_*i*_ and *T* is the duration.

Then, the second step is to work out the total response distribution *P*_*i*_ (*r*), which measures the probability of that the spiking rate of *N*_*i*_ is *r*. In our paper, since the probability distribution of the stimulus 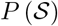 has been given, so we have *P*_*i*_ (*r*) = *P*_*i*_ (*r* | *s*) *P* (*s*).

Finally, we can calculate the parameters respectively as

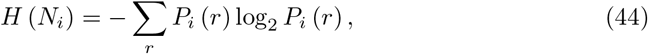

and

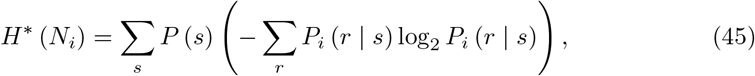

and

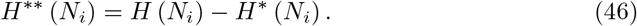

#### Definition of the coding scope

For a given neuron *N*_*i*_ or a neural population {*N*_*i*_}_*i∈I*_ that codes the input stimulus, its efficiently coded information set is defined as

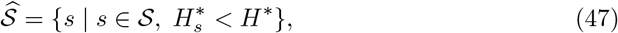

where 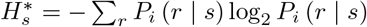 denotes the noise entropy of neural coding for *s* (measures the variation of neural response that can’t be explained by *s*).

Then, we define the coding scope as

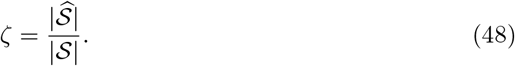

Based on those definitions, it is easy to know that the coding scope *ζ* measures the proportion of the stimuli that can better explain the neural responses.

## Acknowledgments

We would like to thank Annebella Tsz Ho Choi and Weihua He for their assistance in manuscript preparation. This project is supported by Natural Science Foundation of China (81671065) and Tsinghua University Initiative Scientific Research Program.

## Supporting information

**S1. Fig. An example for measuring** *μ*, *θ* **and** ∑ _*j*_ (1 − *τ*_*j*_). In the example, the neural information graph 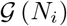 is a complete graph with 4 nodes (meaning that 4 cases of neural information can be transmitted to *N*_*i*_). Although we can directly know that Θ (*N*_*i*_) = 0, we still show the upper bound predicted by our theory is no less than Θ (*N*_*i*_) (of course, we have suggest that the upper bound predicted by us is not supremum when 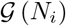 is a complete graph). Given that every node in 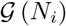 has the same degree (degree is 3) and there is ∑ _*j*_ (1 − *τ*_*j*_) = 0, we can randomly pick one node (rounded by green box) as the node with minimum degree value. Then, we can enumerate all the cliques that contain this node (in gray boxes) and work out that *μ* = 8. Similarly, we enumerate all the cliques in the same connected component with this node and work out that *θ* = 15. Thus, the upper bound predicted by us is given as 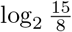. It is clear that the upper bound is greater than Θ (*N*_*i*_) = 0.

**S2 Fig. Several examples of the predicated upper bound by our theory.** (**a-f**) Information rate measurement for the neural information transformation cases with zero-error. All experimental settings remain the same as **Fig. 3c**. (**a-b**) are the experiment for 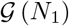, where the upper bound predicted by our theory is log_2_ 3. (**c-d**) are the experiment for 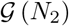, where the upper bound is predicted as log_2_ 7. (**e-f**) are the experiment for 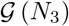, where the upper bound is 2.

**S3 Fig. Several dyeing experiments for neural information confusion detection.**(**a-d**) The dyeing results for various neural populations after encoding the given stimulus sequence in 100 iterations. All experimental settings remain the same as **Fig. 4d**. (**e-f**) The visualization examples of the color variation trajectories for distinguishing confusion and non-confusion. All results are obtained based on the dyeing experiments on a neural population with 1000 neurons.

**S4 Fig. Another example for the effects of neural information confusion on neural coding.** This is another experiment result obtained based on the same settings of the experiment in **Fig. 5**. (**a-b**) The pre-synaptic frequency distributions of *H*, *H*\*, *H*\*\* and *ζ* (averaged between all pre-synaptic neurons), and the corresponding post-synaptic frequency distributions. (**c**) We show the occurrence probability of each variation case and distinguish them based on the variation trend of *H*\*. Then we show the variations of 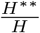 with respect to different cases.

## References

1. Kandel ER, Schwartz JH, Jessell TM, of Biochemistry D, Jessell MBT, Siegelbaum S, et al. Principles of neural science. vol. 4. McGraw-hill New York; 2000.

2. Solso RL, MacLin MK, MacLin OH. Cognitive psychology. Pearson Education New Zealand; 2005.

3. Ghahramani Z. Information theory. Encyclopedia of Cognitive Science. 2006;.

4. Nemenman I, Bialek W, Van Steveninck RDR. Entropy and information in neural spike trains: Progress on the sampling problem. Physical Review E. 2004;69(5):056111.

5. Borst A, Theunissen FE. Information theory and neural coding. Nature neuroscience. 1999;2(11):947.

6. Dayan P, Abbott LF. Theoretical neuroscience: computational and mathematical modeling of neural systems. 2001;.

7. Burkitt AN. A review of the integrate-and-fire neuron model: I. Homogeneous synaptic input. Biological cybernetics. 2006;95(1):1–19.

8. Burkitt AN. A review of the integrate-and-fire neuron model: II. Inhomogeneous synaptic input and network properties. Biological cybernetics. 2006;95(2):97–112.

9. Ochab-Marcinek A, Schmid G, Goychuk I, Hänggi P. Noise-assisted spike propagation in myelinated neurons. Physical Review E. 2009;79(1):011904.

10. Suri RE, Sejnowski TJ. Spike propagation synchronized by temporally asymmetric Hebbian learning. Biological cybernetics. 2002;87(5-6):440–445.

11. Root CM, Semmelhack JL, Wong AM, Flores J, Wang JW. Propagation of olfactory information in Drosophila. Proceedings of the National Academy of Sciences. 2007;104(28):11826–11831.

12. Kumar A, Rotter S, Aertsen A. Spiking activity propagation in neuronal networks: reconciling different perspectives on neural coding. Nature reviews neuroscience. 2010;11(9):615–627.

13. Kish LB, Harmer GP, Abbott D. Information transfer rate of neurons: stochastic resonance of Shannon’s information channel capacity. Fluctuation and Noise Letters. 2001;1(01):L13–L19.

14. Kostal L. Approximate information capacity of the perfect integrate-and-fire neuron using the temporal code. Brain research. 2012;1434:136–141.

15. Suksompong P, Berger T. Capacity analysis for integrate-and-fire neurons with descending action potential thresholds. IEEE Transactions on Information Theory. 2010;56(2):838–851.

16. Bialek W, Zee A. Coding and computation with neural spike trains. Journal of Statistical Physics. 1990;59(1-2):103–115.

17. Schneidman E, Segev I, Tishby N. Information capacity and robustness of stochastic neuron models. In: Advances in neural information processing systems; 2000. p. 178–184.

18. Johnson DH. Information theory and neural information processing. IEEE Transactions on Information Theory. 2010;56(2):653–666.

19. Zador A. Impact of synaptic unreliability on the information transmitted by spiking neurons. Journal of neurophysiology. 1998;79(3):1219–1229.

20. Xu Y, Jia Y, Wang H, Liu Y, Wang P, Zhao Y. Spiking activities in chain neural network driven by channel noise with field coupling. Nonlinear Dynamics. 2019;95(4):3237–3247.

21. White JA, Rubinstein JT, Kay AR. Channel noise in neurons. Trends in neurosciences. 2000;23(3):131–137.

22. Tanabe S, Sato S, Pakdaman K. Response of an ensemble of noisy neuron models to a single input. Physical Review E. 1999;60(6):7235.

23. Shannon C. The zero error capacity of a noisy channel. IRE Transactions on Information Theory. 1956;2(3):8–19.

24. Elias P. Zero error capacity under list decoding. IEEE Transactions on Information Theory. 1988;34(5):1070–1074.

25. Ahlswede R. A note on the existence of the weak capacity for channels with arbitrarily varying channel probability functions and its relation to Shannon’s zero error capacity. The Annals of Mathematical Statistics. 1970;41(3):1027–1033.

26. Lovász L. On the Shannon capacity of a graph. IEEE Transactions on Information theory. 1979;25(1):1–7.

27. Rosenfeld M. On a problem of CE Shannon in graph theory. Proceedings of the American Mathematical Society. 1967;18(2):315–319.

28. Meffin H, Burkitt AN, Grayden DB. An analytical model for the ‘large, fluctuating synaptic conductance state’typical of neocortical neurons in vivo. Journal of computational neuroscience. 2004;16(2):159–175.

29. Butts DA, Goldman MS. Tuning curves, neuronal variability, and sensory coding. PLoS Biol. 2006;4(4):e92.

30. Swindale NV. Orientation tuning curves: empirical description and estimation of parameters. Biological cybernetics. 1998;78(1):45–56.

31. Bosking W, Maunsell J. The correlation between the firing of individual MT neurons and behavioral response across different directions of motion. Soc Neurosci Abs. 2004;31:935–937.

32. Schmitt R, Dev P, Smith BH. Electrotonic processing of information by brain cells. Science. 1976;193(4248):114–120.

33. Cessac B, Paugam-Moisy H, Viéville T. Overview of facts and issues about neural coding by spikes. Journal of Physiology-Paris. 2010;104(1-2):5–18.

34. Stuart G, Spruston N, Sakmann B, Häusser M. Action potential initiation and backpropagation in neurons of the mammalian CNS. Trends in neurosciences. 1997;20(3):125–131.

35. Mazzoni A, Panzeri S, Logothetis NK, Brunel N. Encoding of naturalistic stimuli by local field potential spectra in networks of excitatory and inhibitory neurons. PLoS computational biology. 2008;4(12).

36. Simoncelli EP, Heeger DJ. A model of neuronal responses in visual area MT. Vision research. 1998;38(5):743–761.

37. Simoncelli E, Bair W, Cavanaugh J, Movshon JA. Testing and refining a computational model of neural responses in area MT. Investigative Ophthalmology and Visual Science. 1996;37(3).

38. Simoncelli E, Heeger D. A velocity representation model for MT cells. Investigative Opthamology and Visual Science Supplement. 1994;35:1827.

39. Averbeck BB, Latham PE, Pouget A. Neural correlations, population coding and computation. Nature reviews neuroscience. 2006;7(5):358.

40. Oram MW, Földiák P, Perrett DI, Sengpiel F. TheIdeal Homunculus’: decoding neural population signals. Trends in neurosciences. 1998;21(6):259–265.

41. Panzeri S, Schultz SR, Treves A, Rolls ET. Correlations and the encoding of information in the nervous system. Proceedings of the Royal Society of London Series B: Biological Sciences. 1999;266(1423):1001–1012.

42. Shadlen MN, Newsome WT. The variable discharge of cortical neurons: implications for connectivity, computation, and information coding. Journal of neuroscience. 1998;18(10):3870–3896.

43. Moreno-Bote R, Beck J, Kanitscheider I, Pitkow X, Latham P, Pouget A. Information-limiting correlations. Nature neuroscience. 2014;17(10):1410.

44. Romo R, Hernández A, Zainos A, Salinas E. Correlated neuronal discharges that increase coding efficiency during perceptual discrimination. Neuron. 2003;38(4):649–657.

45. Kanitscheider I, Coen-Cagli R, Pouget A. Origin of information-limiting noise correlations. Proceedings of the National Academy of Sciences. 2015;112(50):E6973–E6982.

